# One-Pot Time-Induced Proteome Integral Solubility Alteration (OPTI-PISA) assay for automated and sensitive drug target identification

**DOI:** 10.1101/2024.08.13.607299

**Authors:** Zhaowei Meng, Amir Ata Saei, Hezheng Lyu, Massimiliano Gaetani, Roman A. Zubarev

## Abstract

Proteome Integral Solubility Alteration (PISA) assay is widely used for identifying drug targets, but it is labour-intensive and time-consuming, and requires a substantial amount of biological sample. Aiming at enabling automation and greatly reducing the sample amount, we developed One-Pot Time-Induced (OPTI)-PISA. Here we demonstrate OPTI-PISA performance on identifying targets of multiple drugs in cell lysate and scaling down the sample amount to sub-microgram levels, making PISA method suitable for NanoProteomics.

---

As proteins are cellular building blocks and indispensable elements of cell mechanics, most processes that occur in a cell or organism involve proteins^1-3^. Protein higher-order structures undergo changes when proteins in a cell, bodily fluid or in an isolated state interact with other molecules, including proteins^4,5^, metabolites^6,7^ and drug molecules^8^. Monitoring these changes in protein structure is important to understand how cells live, metabolise food, proliferate, communicate with each other, mutate, differentiate and die^9-12^.

In biology and medicine, it is often important to identify which protein in a whole proteome interacts with an active molecule (e.g., a drug). Thermal Proteome Profiling (TPP^13^), also known as CETSA-MS^8^, is a method for identifying drug targets by measuring changes in the thermal stability of proteins in a proteome-wide manner. Protein thermal stability is probed by incubating proteome solution at a set of temperature points varying from normal to elevated (e.g., 37 °C to 67 °C), centrifugation and performing proteomic analysis of the supernatant at each temperature. At elevated temperatures, native conformations of many proteins unfold (‘melt’), which makes most proteins less soluble and ultimately results in protein aggregation. TPP measures the shift in protein melting temperature (T_m_), which is postulated to occur when the protein binds a ligand. A typical TPP workflow involves subjecting samples to ∼10 different temperature points and using one tandem mass tag (TMT) set per sample to obtain T_m_ values by fitting a sigmoidal curve into relative protein abundance at all temperature points. To establish the ligand-induced shift in T_m_, this procedure needs to be performed separately for drug-treated and control samples^14^, and in order to assess statistical significance of the results, this has to be done in a number of replicates. TPP has been used quite extensively^15-18^, but it is plagued by a limited throughput, large sample requirements and high labour intensity. Another disadvantage of TPP is that many melting curves fit poorly to sigmoidal shape, which prevents precise T_m_ measurements^14,19^.

Proteome integral solubility alteration (PISA)^20^ has been developed as a high-throughput alternative to TPP^5,21-23^. In PISA, the samples heated to different temperatures are combined, and the overall change in protein solubility (which is equivalent to the area under the melting curve in TPP) is measured. PISA increases the analysis throughput by 1 to 2 orders of magnitude and significantly reduces the required sample amount compared to TPP^24^. However, classical PISA is still not capable of analyzing low-microgram amounts of samples, because dividing the sample into multiple vessels for thermal treatment results in significant sample loss on the walls of the tubes and pipettes. Moreover, PISA largely inherited from TPP the manual labour intensity. Automation of PISA could greatly reduce the labour demand and increase the repeatability of the results. A more precise analysis would require fewer replicates to obtain the desired statistical power, which would result in significant reduction of the analysis time and sample consumed.

Automation of PISA requires solving the problem of heating different sample portions to different temperatures, and then combining these portions in a single volume. Initially we solved this problem by injecting the sample of lysate with or without the drug into a straight capillary (‘one pot’) attached to a metal plate. The plate was heated by two heaters at both ends such that a temperature gradient existed across the plate and thus across the capillary. The injected sample stayed in a differentially heated capillary for a few minutes, and then was eluted to a vial and centrifuged. Different parts of the same sample were thus exposed to different temperatures without the need of keeping them in separate vessels. This approach also obviates the need for splitting the sample into different tubes for incubation at different temperatures. The approach was tested and found workable, showing potential to be automated. However, maintaining precise temperature gradient turned out to be a challenge during long runs. In a modern laboratory, the ambient air temperature is controlled by a computer that can be programmed to reduce it at night and over the weekends. As the temperature gradient on the heated PISA plate depends upon the ambient conditions, changes in air temperature may affect that gradient, resulting in experiment-to-experiment variations.

Thus, we chose an alternative approach, using the fact that protein solubility can be modulated not only by heating sample to different temperatures for the same time duration, as in TPP and PISA, but also by heating different sample volumes to the same temperature but for different time intervals. This is somewhat similar to egg boiling, in which different boiling times result in different softness of the eggs. In that approach termed One-Pot Time-Induced (OPTI)-PISA, the metal plate is uniformly heated to an intermediate temperature (which is much easier to maintain constant than a temperature gradient) at which different parts of the sample in a capillary are incubated for different time intervals. The latter is achieved by a fast injection of the sample into the heated capillary and a slower elution from it (**Fig. 1**). Automated injection and elution ensure repeatable experimental conditions consistently applied to different samples, which reduces the statistical uncertainty of the results compared to manual sample handling. Also, performing incubation in a single vessel enables dramatic reduction in sample volume.

**Fig. 1.**
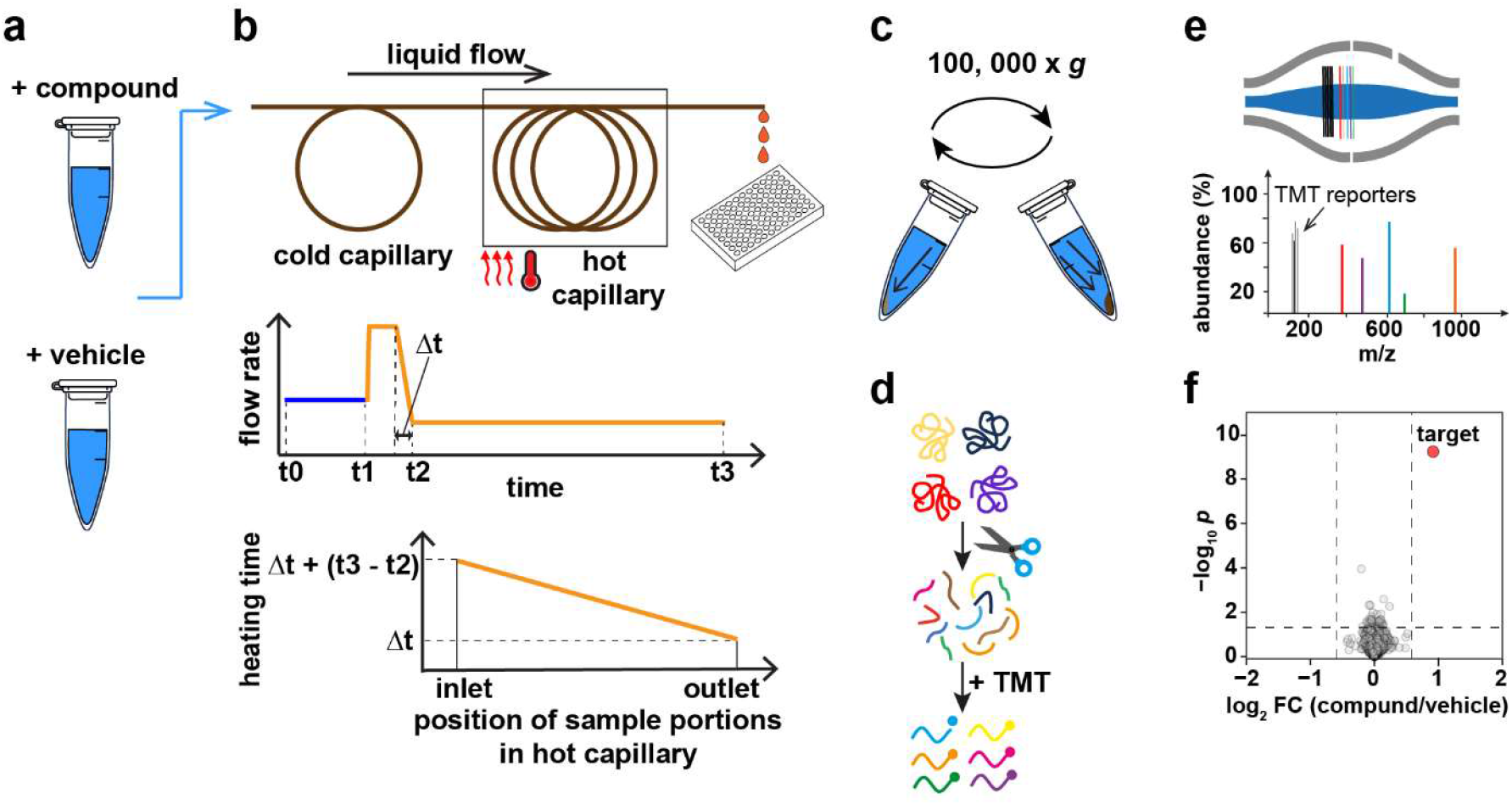
Schematic diagram of the OPTI-PISA workflow. (**a**) The cell lysate is treated with drug or vehicle. (**b**, upper). The treated sample is first injected into the cold sample loop and then transferred to a heated capillary kept at a constant temperature, and after a short time interval eluted at a slow flow rate. (**b**, middle) The flow rate through the capillary. (**b**, lower) Exposure time of sample molecules located in different capillary parts to elevated temperature. (**c**) Ultra-centrifugation and removal of aggregated proteins. (**d**) Proteomic sample preparation. (**e**) LC-MS/MS analysis. (**f**) Data processing and analysis.

In the proof of principle of the OPTI-PISA assay, we employed an automated liquid handling system equipped with a pump, autosampler and a fractionator (Biomotif) (**Supplementary Fig. 1**). A549 cell lysate (17.5 μg/sample) with and without addition of 10 μM methotrexate (MTX) was used as a model system. The results were compared between the manually controlled injection/elution and fully automatic sample processing. The plate was heated to 50 °C, the injection rate was 50 μL/min, while the elution rate was 100 times slower, 0.5 μL/min. All samples were processed in five replicates multiplexed into a single TMT-10 labeling set. In both manual and automated processing, the subsequent LC-MS/MS analysis correctly identified DHFR as a sole target among 3722 and 3291 proteins, respectively, quantified with ≥ 2 unique peptides (**Fig. 2a-b**). As expected, OPTI-PISA automation improved the p value from 7 × 10^-6^ for the manual setup to 5.8 × 10^-10^ in automated sample processing. In terms of statistical significance, this improvement was equivalent to roughly doubling the number of replicates in manual analysis.

**Fig. 2.**
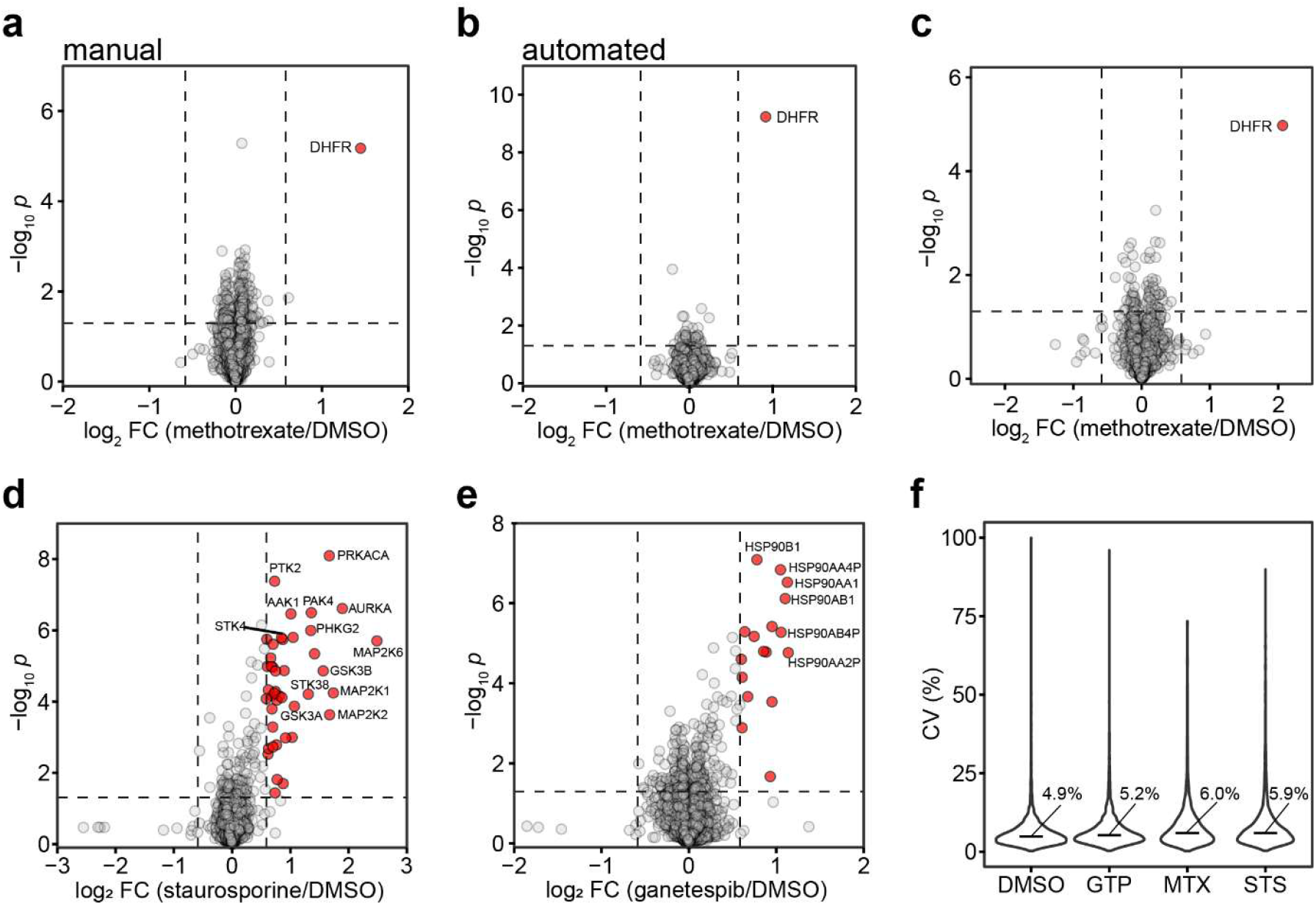
OPTI-PISA results for A549 cell lysate incubated with 10 μM drugs, versus vehicle incubation. (**a-c**) lysate incubated for 1 h with drugs. (**a**) MTX, 17.5 μg of lysate per sample, manual injection/elution. (**b-f**) Automated OPTI-PISA. (**b**) MTX, 17.5 μg of lysate per sample (**c**) MTX, 900 ng of lysate per sample. (**d**) Staurosporine, 150 μg of lysate per sample, incubated for 15 min. (**e**) Ganetespib, 150 μg of lysate per sample, incubated for 30 min. (**f**) CVs of protein abundances. Median values are shown.

To test the suitability of OPTI-PISA for NanoProteomics, the protein amount was reduced to 900 ng per replicate. DHFR was still identified as the sole MTX target (**Fig. 2c**) among the 1443 proteins quantified with > 2 unique peptides.

To generalize the OPTI-PISA approach, we tested its performance on a standard HPLC set-up equipped with an autosampler and fraction collector (**Supplementary Fig. 2**). The latter were used for liquid handling, while an external heater was used for heating of the sample inside the capillary attached to the heater. After thermal treatment, each sample was frozen in liquid nitrogen before processing. As a test system, A549 lysate was treated with 3 drugs at a 10 μM concentration and a vehicle as a control, with 4 replicates of each treatment multiplexed into a single TMT-16 set. As drugs we used staurosporine (a broad spectrum protein kinase inhibitor ^25^), ganetespib (a HSP90 inhibitor^26,27^) and MTX (**Supplementary Fig. 3a**). In total, 5408 proteins were quantified with ≥ 2 unique peptides. For staurosporine, 38 kinases showed a significant increase in solubility (**Supplementary Fig. 3b**). Of these, 28 (74%) were among the 268 known targets of staurosporine^28^, while random overlap would give on average only 2 common proteins. For ganetespib, HSP90 had the most significant increase in solubility (**Supplementary Fig. 3c)**, in agreement with literature^26^. Importantly, all median coefficients of variability (CVs) of the protein abundances across the replicates were lower than 8% (**Supplementary Fig. 3d**). This result highlighted excellent reproducibility of this OPTI-PISA set-up, even though pipetting of the cell lysate into individual sample wells and adding drugs/vehicle to each sample were still manually performed.

To make OPTI-PISA completely automated, we employed the HPLC column oven for thermal treatment, and programmed the autosampler to automatically pipette cell lysate and drugs/vehicle from their storage vial, while the fraction collector was instructed to store the thermally treated samples at 4 °C before further processing. The performance of completely automated OPTI-PISA set-up was done using the same test system as above. In total, 6184 proteins were quantified with ≥ 2 unique peptides. The positive control, DHFR protein, was identified as a sole target of MTX (**Supplementary Fig. 4a)**. The staurosporine results revealed 44 kinases with a significant increase in solubility **(Fig. 2d)**, 31 (70%) of which were among the 268 known targets of staurosporine (random overlap would produce only 2 common proteins). In ranking the quantified proteins by their log2-scaled fold change multiplied by −log10-scaled p value, the top 40 proteins were all kinases (**Supplementary Fig. 4b)**. For ganetespib, HSP90 exhibited the most significant increase in solubility **(Fig. 2e)**, while the top 6 proteins in the above ranking were all proteoforms of HSP90 (**Supplementary Fig. 4c)**. The median CVs of all protein abundances across the 4 replicates were lower than 7% **(Fig. 2f)**, highlighting the superior reproducibility of completely automated OPTI-PISA.

In conclusion, the OPTI-PISA approach afforded easy and complete automation of sample thermal treatment in PISA experiments, offering higher sensitivity and repeatability, with lower sample and labour requirements.

## METHODS

### Cell Culture

Human lung carcinoma A549 cells (ATCC) were grown at 37°C in 5% CO_2_ using Dulbecco’s Modified Eagle Medium (Lonza) supplemented with 10% FBS superior (Biochrom) and 100 units/mL penicillin/streptomycin (Gibco). Low-number passages (<10) were used for the experiments. Cellular lysates were obtained in PBS by five cycles of freezing cells in liquid N_2_ and then thawing at 25 °C, then cleared by centrifugation at 14,000 rpm for 10 min at 4 °C. The protein concentration in the lysate was measured using Pierce BCA kit (Thermo Fisher).

### Manual OPTI-PISA Configuration for MTX experiment

The device consists of a syringe pump (Chemyx), 100 μL syringes (Hamilton), connected by a union to fused silica capillary (75 μm × 113 cm, 5 μL volume) attached to a metal plate kept at a constant temperature (50 °C) by a programmable heater. The sample fills the capillary at a flow rate of 100 μL/min for 10 s, then the syringe pump stops the flow for 10 s. Subsequently, the syringe pump elutes the sample at a flow rate of 0.5 μL/min for 10 min, and the elution is collected into a 0.5 mL-LoBind tube (Eppendorf) containing 95 μL phosphate buffered saline (PBS). The syringe and capillary are washed with PBS before running the next sample.

### Automated OPTI-PISA Configuration for MTX experiments

In the manual OPTI-PISA instrument, the syringe pump is replaced by a programmable liquid sample handler comprising a pump and an autosampler with a fractionation collection option. 15 μL of sample is drawn up from a 96-well plate into a 10 μL sample loop. The pump pushes PBS at 20 μL/min to wash capillary for 2 min, and fast transfers the sample from the sample loop into the heated capillary at a flow rate of 50 μL/min for 8 s, then stops for 10 s for stabilizing the flow. Then the pump elutes the sample from the hot capillary into the 96-well plate at a flow rate of 0.5 μL/min for 5 min, and then switches to another well for 10 min. The aim is to collect two fractions per sample, one (≈5 μL) subjected to intended thermal treatment that will be used for analysis, and another one wasted as an overhead.

### Automated OPTI-PISA Configuration for Multiple Drug experiment

Ultimate 3000 HPLC system with fractionation option (Thermo Fisher) was employed. As an external device, a fused silica capillary (250 μm × 102 cm, 50 μL volume) was placed on a metal plate and heated to 55 °C by a programmable thermal heater. The autosampler picked 70 μL of sample into a 125 μL sample loop, and the pump fast transferred sample from sample loop to the heating capillary at a flow rate of 200 μL/min for 15 s, then reduced the flow rate to 10 μL/min to elute the sample from the hot capillary into a 96-well plate.

### Completely Automated OPTI-PISA Configuration for Multiple Drug experiment

Ultimate 3000 HPLC system with fractionation option (Thermo Fisher) was employed. A fused silica capillary (380 μm × 100 cm, 100 μL volume) was placed in HPLC column oven heated to 55 °C. The autosampler picked 100 μL of sample into a 125 μL sample loop, and the pump fast transferred sample from sample loop to the heating capillary at a flow rate of 200 μL/min for 15 s, then reduced the flow rate to 20 μL/min to elute the sample from the hot capillary into a 96-well plate.

### OPTI-PISA Experiments with MTX

A549 cellular lysate was treated with MTX at 10 μM final concentration or an equal amount of DMSO (control) with 5 replicates in PBS for 60 min at 25°C. Samples were respectively subjected to thermal treatment on the manual OPTI-PISA configuration or automated OPTI-PISA instrument as described above. Collected sample elution was then left at room temperature for 4 min before being snap frozen in liquid N_2_. The samples were then transferred into polycarbonate thick-wall tubes and centrifuged at 100,000 g at 4 °C for 30 min. The supernatant was collected. Lys-C (Wako) was added at a 1:40 w/w ratio and samples were incubated at 30 °C overnight. Sequencing grade modified trypsin (Promega) was then added at a 1:20 w/w ratio, and the samples were digested at 37 °C for 4 h. TMT 10plex reagents dissolved in acetonitrile (ACN) were added 4× by weight to each sample with a final ACN concentration of 30%, followed by incubation for 2 h at 25 °C. The reaction was quenched by adding 0.5% hydroxylamine. Samples were combined, acidified by TFA to pH 2-3. Sample desalting and fractionation was performed by high-pH reverse-phase fractionation Kit (Thermo Fisher). A total of 12 fractions were collected and concatenated into 10 fractions. Sub-microgram samples were cleaned by StageTips without fractionation after TMT-10 labelling. Samples were dried in a DNA 120 SpeedVac Concentrator (Thermo Fisher) and stored at −80 °C until LC-MS/MS analysis.

### OPTI-PISA Experiment with Multi-target Drugs

3 drugs (methotrexate, staurosporine, ganetespib) at a final concentration of 10 μM and DMSO (control) were used for treating the A549 lysate in 4 replicates. The A549 lysate (1.5 μg/μL) was distributed into 16 equal aliquots in sample vials (80 μL/vial) and these vials were stored at 4 °C. 1 μL of 800 μM drug or same concentration of DMSO was added to each sample vial containing 80 μL of A549 lysate, pipetting 10 times without making bubbles. After incubating at 25 °C for 15 min, the sample was deposited in the autosampler of Ultimate 3000 HPLC system for OPTI-PISA experiment as described above. The eluted sample was left at room temperature for 4 min before being snap frozen in liquid N_2_. Samples were then transferred into polycarbonate thick-wall tubes and centrifuged at 100,000 × g for 30 min at 4 °C. The supernatant was collected, and its protein concentration was measured using Pierce BCA kit. Dithiothreitol (DTT) was added to a final concentration of 8 mM and samples were incubated for 45 min at 55 °C. Subsequently, iodoacetamide (IAA) was added to a final concentration of 25 mM and samples were incubated at 25 °C for 30 min in the darkness. Proteins were precipitated using cold acetone at a sample: acetone ratio of 1:6 (v/v) at −20 °C overnight. The precipitated protein was collected by centrifugation (10,000 × g, 10 min), and the supernatant was removed. The protein pellet was air-dried for 3 min, then dissolved in 10 μL of 20 mM EPPS buffer at pH 8.2 containing 8 M urea for 10 min. 20 mM EPPS buffer (pH 8.2) was added to dilute urea to 4M. Then 1 μL of LysC was added at a 1:75 w/w ratio, the samples were incubated at 30 °C for 6 h. The samples were diluted with 20 mM EPPS to a final urea concentration of 1 M, and trypsin was added at a 1:50 w/w ratio, followed by incubation at 37 °C overnight. TMTpro 16plex reagents dissolved in acetonitrile (ACN) were added 5× by weight to each sample with a final ACN concentration of 30%, followed by incubation for 2 h at 25 °C. The reaction was quenched by adding 0.5% hydroxylamine. Samples were combined, acidified by TFA to pH 1-3, cleaned using Sep-Pak cartridges (Waters), and dried using DNA 120 SpeedVac Concentrator (Thermo Fisher). Peptide separation for deeper proteome analysis was carried out using an Ultimate 3000 HPLC system (Thermo Fisher) equipped with a Xbridge Peptide BEH C18 column (25 cm × 2.1 mm, particle size 3.5 μm, pore size 300 Å; Waters) operating at a flow rate of 200 μL/min. Fractionation was achieved through a binary solvent system comprising 20 mM NH_4_OH in H_2_O (solvent A) and 20 mM NH_4_OH in acetonitrile (solvent B). The elution profile was programmed as follows (data for % of solvent B): 2% to 23% in 42 min, 23 to 52% in 4 min, 52 to 63% in 2 min, and a subsequent isocratic hold at 63% for 5 min. The elution process was monitored by continuous of UV absorbance at 214 nm. A total of 96 fractions, each containing a 100 μL aliquot, were collected. Fractions were then concatenated into 24 samples in a sequential order (e.g. 1, 25, 49, 73).

### Completely Automated OPTI-PISA Experiment with Multi-target Drugs

An Ultimate 3000 HPLC system was employed to perform all the processes in an automatic manner. 3 drugs (methotrexate, staurosporine, ganetespib) at a final concentration of 10 μM and DMSO (control) were used for treating the A549 lysate in 4 replicates. The A549 lysate (1.5 μg/μL) was automatically distributed into 16 equal aliquots in the wells of a 96-well plate (108 μL/well) and these vials were stored at 4 °C. 12 μL of 100 μM drug or same concentration of DMSO was automatically added to each sample well containing 108 μL of A549 lysate, shaking by the autosampler tray without making bubbles. After incubating at 25 °C for 30 min, the sample was processed for completely automated OPTI-PISA experiment as described above. Samples were then transferred into polycarbonate thick-wall tubes and centrifuged at 100,000 g for 30 min at 4 °C. The proteomics sample preparation was the same as described in **OPTI-PISA Experiment with Multi-target Drugs**. After high-pH fractionation, fractions were then concatenated into 48 samples in a sequential order (e.g. 1 and 49, 2 and 50).

### LC-MS/MS Analysis

Each sample was analysed by LC-MS/MS using a Q Exactive HF or Orbitrap Exploris 480 mass spectrometers equipped with an EASY Spray Ion Source and connected to an Ultimate 3000 RSLC nano HPLC system (all - Thermo Fisher). Injected sample fractions were preconcentrated and further desalted on a trap column (Acclaim PepMap 100 C18, 75 μm × 2 cm; particle size, 3 μm; pore size, 100 Å), and then separated on an EASY Spray analytical column (Acclaim PepMap RSLC, 50 cm × 75 μm; particle size, 2 μm; pore size, 100 Å) at 55°C and a flow rate of 300 nL/min. Peptides were separated using a binary solvent system consisting of 0.1% (v/v) formic acid (FA), 2% (v/v) acetonitrile (ACN) (solvent A) and 98% ACN (v/v), 0.1% (v/v) FA (solvent B).

For the fractionated samples, the gradient consisted of 3−28% B in 115 min, 28−40% B in 5 min, 40−95% B in 5 min, 95% B for 8 min, and 95-3% B in 2 min. For the unfractionated sub-microgram sample, the gradient consisted of 3−28% B in 230 min, 28−40% B in 10 min, 40−95% B in 5 min, 95% B for 10 min, and 95-3% B in 2 min.

For the fractionated samples from OPTI-PISA experiment with single-target drug (MTX), mass spectra were acquired using Q Exactive HF. A full MS (MS1) spectrum was first acquired in the Orbitrap analyzer with mass-to-charge ratio (m/z) range from 375 to 1500, nominal resolution 120,000, automated gain control (AGC) target 3 × 10^6^, and maximum injection time of 100 ms. Following this, the 17 most abundant peptide ions were selected for higher-energy collision dissociation (HCD) with normalized collision energy 33 for subsequent MS/MS (MS2) analysis with a minimum intensity threshold of 2 × 10^4^ and a 30 s dynamic exclusion time. MS2 spectra were acquired in the Orbitrap analyzer with the following settings: quadrupole isolation window 1.2 Th, AGC target 2 × 10^5^, maximum injection time 120 ms, nominal resolution 45,000, and fixed first m/z 110.

For the fractionated samples obtained from OPTI-PISA assay for 3 drugs, mass spectra were acquired using Orbitrap Exploris 480. A full MS (MS1) spectrum was first acquired with m/z range from 375 to 1500, nominal resolution 120,000, AGC target 3 × 10^6^, and maximum injection time 50 ms. The most abundant peptide ions were selected for HCD with normalized collision energy 35 for subsequent MS/MS (MS2) analysis with a minimum intensity threshold of 5 × 10^4^ and a 30 s dynamic exclusion time. MS2 spectra were acquired at nominal resolution 45,000, with AGC target 2.5 × 10^5^ and maximum injection time 120 ms. The fixed first m/z was 110 and the isolation window was 0.7 Th. The number of MS2 spectra acquired per each MS1 spectrum was determined by setting the maximum cycle time for MS1 and MS2 spectra to 3 s (using the top speed mode).

For the unfractionated sub-microgram sample, Orbitrap Exploris 480 was used for mass spectra acquisition. All the parameters were the same as used for the fractionated samples obtained from OPTI-PISA with 3 drugs, except for normalized collision energy set to 33 and dynamic exclusion time 60 s.

### Data Processing

The raw LC-MS/MS data were analysed by MaxQuant, version 2.2.0.0. The Andromeda search engine was employed to perform MS/MS data matching against the UniProt Human proteome database (version UP000005640_9606, 20607 human protein sequences). Enzyme specificity was set to trypsin, with maximum two missed cleavages permitted. Cysteine carbamidomethylation was set as a fixed modification, while methionine oxidation, N-terminal acetylation, asparagine or glutamine deamidation were used as variable modifications. 1% false discovery rate was used as a filter at both protein and peptide levels. Default settings were employed for all other parameters. Peptide quantification was executed using the abundances of reporter ions of TMT 10 plex or TMTpro 16plex according to the experiments. The obtained protein abundances were normalized on the total ion abundance of TMT reporters for a given protein, then the average value of protein abundance across replicates was calculated. Volcano plots were generated by calculating the protein abundance fold change (FC) between drug and DMSO treated samples and performing Student’s t-tests for the difference between the two treatment groups.

### Accession codes

The raw mass spectrometric data used in this study are available at ProteomeXchange with the identifier PXD050241.

## ACKNOWLEDGEMENTS

We are grateful to Marie Ståhlberg and Carina Palmberg for their assistance in proteomic sample preparation and Hassan Gharibi for improving figure quality.

## AUTHOR CONTRIBUTIONS

Conceptualization, R.A.Z.; methodology and experiment design, R.A.Z., Z.M., A.A.S.; experiments, Z.M., A.A.S., H.L. and M.G.; project organization, training, resources and funding acquisition, R.A.Z., M.G.; data analysis and visualization, Z.M., A.A.S.; manuscript writing R.A.Z., Z.M.

