## Supplementary Figures for "One-Pot Time-Induced Proteome Integral Solubility Alteration (OPTI-PISA) assay for automated and sensitive drug target identification"

Roman A. Zubarev[1,2,3*]

[1] Division of Chemistry I, Department of Medical Biochemistry and Biophysics, Karolinska Institutet, 17177 Stockholm, Sweden

[2] Chemical Proteomics Unit, Science for Life Laboratory (SciLifeLab), 17165 Stockholm, Sweden

[3] Chemical Proteomics, Swedish National Infrastructure for Biological Mass Spectrometry (BioMS), 17177 Stockholm, Sweden

[4] Department of Microbiology, Tumor and Cell Biology, Karolinska Institutet, 17177 Stockholm, Sweden

[5] HDXperts AB, 18212 Danderyd, Sweden

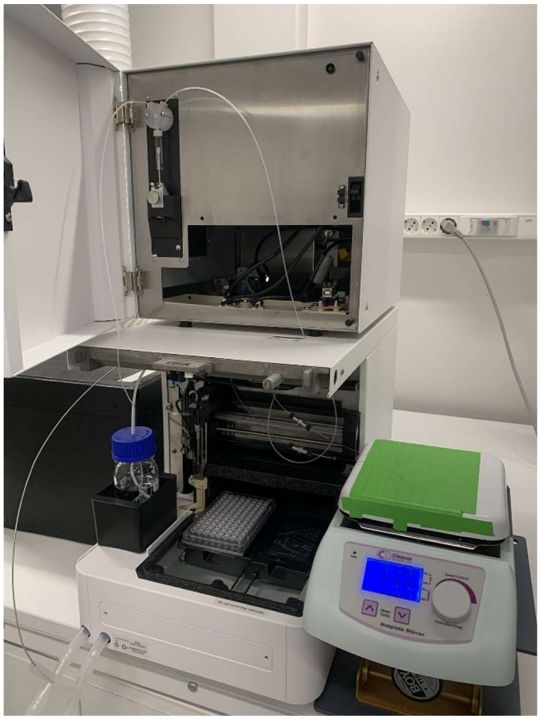

**Supplementary Fig. 1. An automated liquid handling system for proof-of-principle OPTI-PISA experiments.** This system is equipped with a pump, autosampler and fractionator (Biomotif, Danderyd, Sweden).

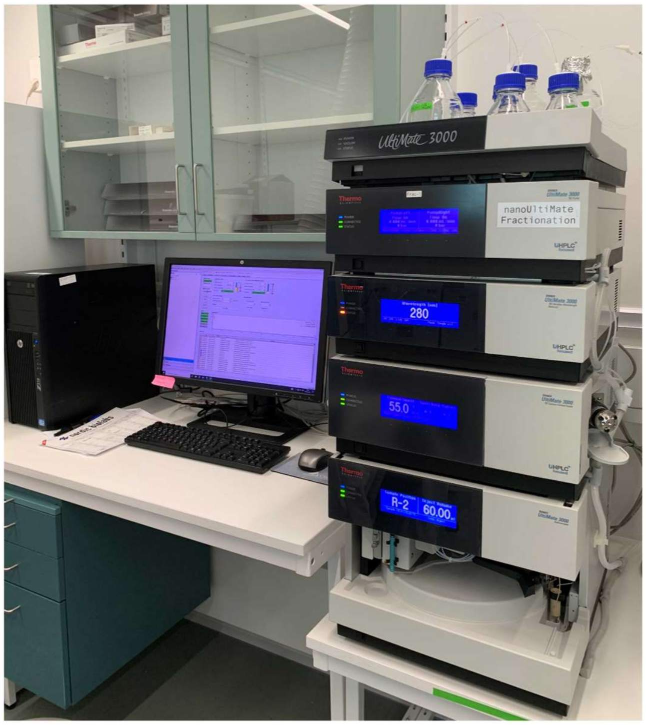


**Supplementary Fig. 2. A standard HPLC system for OPTI-PISA experiments.** This system is a widely used Thermo Ultimate 3000 HPLC equipped with an autosampler combined with fraction collector.

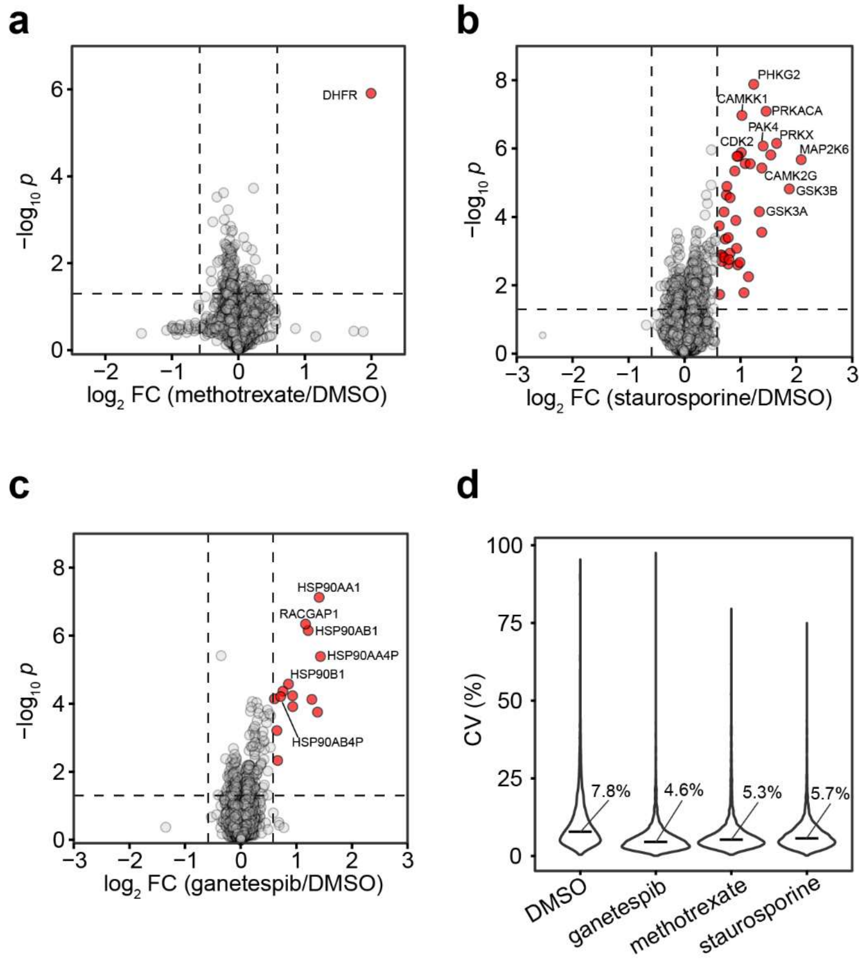


**Supplementary Fig. 3. Automated OPTI-PISA results for A549 cell lysate and 10 μM drugs.** (**a**) MTX. (**b**) Staurosporine. (**c**) Ganetespib. (**d**) CVs of protein abundances. (**a-c**) 70 μg/sample.

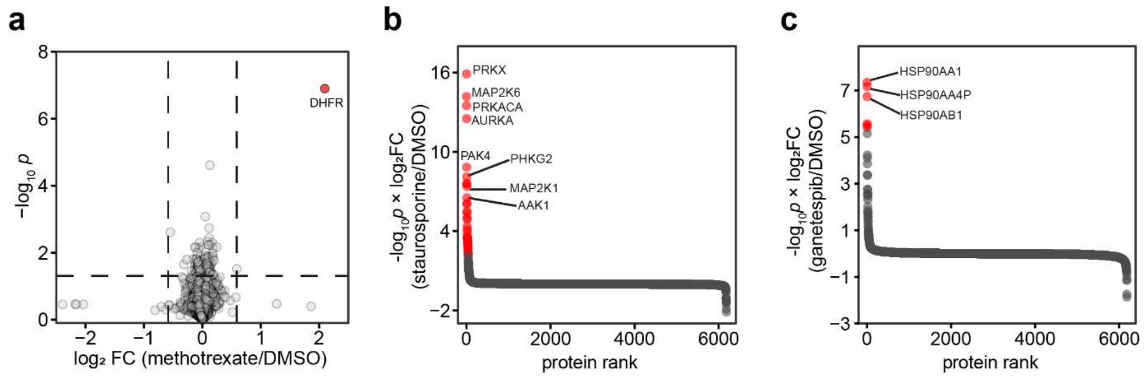


**Supplementary Fig. 4. Results from fully automated OPTI-PISA for A549 cell lysate and 10 μM drugs.** (**a**) MTX as a positive control. (**b**) Ranks of quantified proteins sorted by their log2-scaled fold change multiplied by -log10-scaled p value for staurosporine and (**c**) ganetespib.
